# Bioorthogonal labeling and enrichment of histone monoaminylation reveal its accumulation and regulatory function in cancer cell chromatin

**DOI:** 10.1101/2024.03.20.586010

**Authors:** Nan Zhang, Jinghua Wu, Farzana Hossain, Haidong Peng, Huapeng Li, Connor Gibson, Min Chen, Huan Zhang, Shuaixin Gao, Xinru Zheng, Yongdong Wang, Jiangjiang Zhu, Jing J. Wang, Ian Maze, Qingfei Zheng

**Affiliations:** Department of Radiation Oncology, College of Medicine, The Ohio State University, Columbus, OH 43210, United States; Center for Cancer Metabolism, James Comprehensive Cancer Center, The Ohio State University, Columbus, OH 43210, United States; Molecular, Cellular, and Developmental Biology Graduate Program, The Ohio State University, Columbus, OH 43210, United States; Nash Family Department of Neuroscience, Friedman Brain Institute, Icahn School of Medicine at Mount Sinai, New York, NY 10029, United States; Human Nutrition Program, Department of Human Sciences, College of Education and Human Ecology, The Ohio State University, Columbus, OH 43210, United States; Department of Cancer Biology and Genetics, Comprehensive Cancer Center, The Ohio State University, Columbus, OH 43210, United States; Cerno Bioscience, Las Vegas, NV 89144, United States; Department of Pharmacological Sciences, Icahn School of Medicine at Mount Sinai, New York, NY 10029, United States; Howard Hughes Medical Institute, Icahn School of Medicine at Mount Sinai, New York, NY 10029, United States; Department of Biological Chemistry and Pharmacology, College of Medicine, The Ohio State University, Columbus, OH 43210, United States

**Author notes:** To whom correspondence should be addressed: Prof. Qingfei Zheng, Tzagournis Medical Research Facility, 420 W. 12th Ave. Columbus, OH 43210, United States. These authors contributed equally (Nan Zhang, Jinghua Wu, Farzana Hossain).

## Abstract

Histone monoaminylation (*i*.*e*., serotonylation and dopaminylation) is an emerging category of epigenetic mark occurring on the fifth glutamine (Q5) residue of H3 N-terminal tail, which plays significant roles in gene transcription. Current analysis of histone monoaminylation is mainly based on site-specific antibodies and mass spectrometry, which either lacks high resolution or is time-consuming. In this study, we report the development of chemical probes for bioorthogonal labeling and enrichment of histone serotonylation and dopaminylation. These probes were successfully applied for the monoaminylation analysis of *in vitro* biochemical assays, cells, and tissue samples. The enrichment of monoaminylated histones by the probes further confirmed the crosstalk between H3Q5 monoaminylation and H3K4 methylation. Finally, combining the *ex vivo* and *in vitro* analyses based on the developed probes, we have shown that both histone serotonylation and dopaminylation are highly enriched in tumor tissues that overexpress transglutaminase 2 (TGM2) and regulate the three-dimensional architecture of cellular chromatin.

**TOC:** 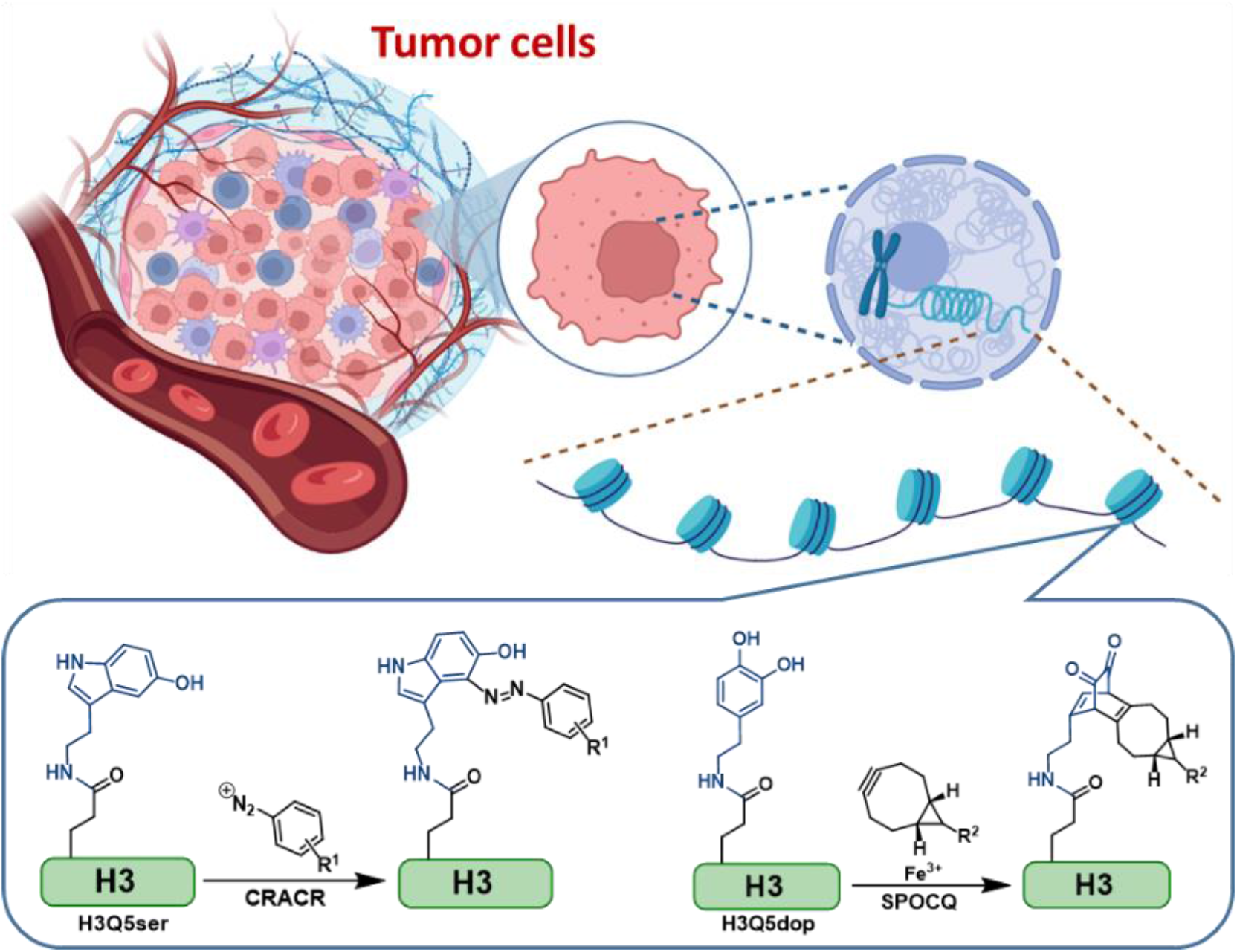

---

The hereditary material of eukaryotic cells is packaged in the nucleus as a dynamic nucleoprotein chromatin complex, where an octamer of core histones (*i*.*e*., two copies of H2A, H2B, H3, and H4) is spooled around 1.5 times by 147 base pairs of DNA to form the basic repeating unit of chromatin organization, nucleosome.^1^ Previous studies have implied that post-translational modifications (PTMs) on histones (also known as the “histone code”),^2^ which are dynamically regulated by “writer” and “eraser” enzymes and recognized by “reader” proteins, are a category of essential epigenetic modifications playing a role in both health and disease states.^3^ The dynamic installation, removal, and recognition of histone PTMs (*e*.*g*., methylation, acetylation, and phosphorylation) have been proven to directly affect cellular phenotypes through manipulating gene transcription and chromatin structures.^4^ These pathways also serve as potential targets for human disease treatment, including cancer.^5,6^

In our previous studies, we reported a series of cellular microenvironment-driven histone PTMs as novel epigenetic markers that are induced by chemically reactive metabolites within the tumor microenvironment (TME).^7,8^ These modifications, which have direct impacts on gene transcription and chromatin three-dimensional architectures, include non-enzymatic glycation caused by reducing sugar molecules, such as methylglyoxal (MGO),^9^ glyoxal,^10^ ribose,^11^ and glucose.^12^ The higher glycolytic rate and enrichment of MGO in tumor cells make histone MGO-glycation a hallmark of cancer.^9,13,14^ Recently, we elucidated the biochemical mechanism of another type of intriguing microenvironment-driven histone PTM, *i*.*e*., monoaminylation on N-terminal glutamine residue of histone H3 (H3Q5).^15,16,17^ The dynamics of this modification, including its installation, removal, and replacement, are regulated by a single enzyme, transglutaminase 2 (TGM2) through a transamination reaction (**Fig. 1A**).^17^ In addition to its unique biochemical mechanism, histone monoaminylation is essential in gene regulation. It has been demonstrated that H3 monoaminylation plays significant roles in regulating neuronal transcription, both during development and in the adult brain.^15,16^ Recently, we discovered that histone monoaminylation was able to promote neural rhythmicity through epigenetic regulations.^17^ Therefore, to detect the histone monoaminylation levels in different types of cells and tissues is of great interest to researchers for both basic and translational studies (such as disease diagnosis).^18^ However, due to the insensitivity and nonspecificity of the corresponding antibodies, the precise and rapid detection of histone monoaminylation in tissue or cell samples still remains challenging.

**Figure 1.**
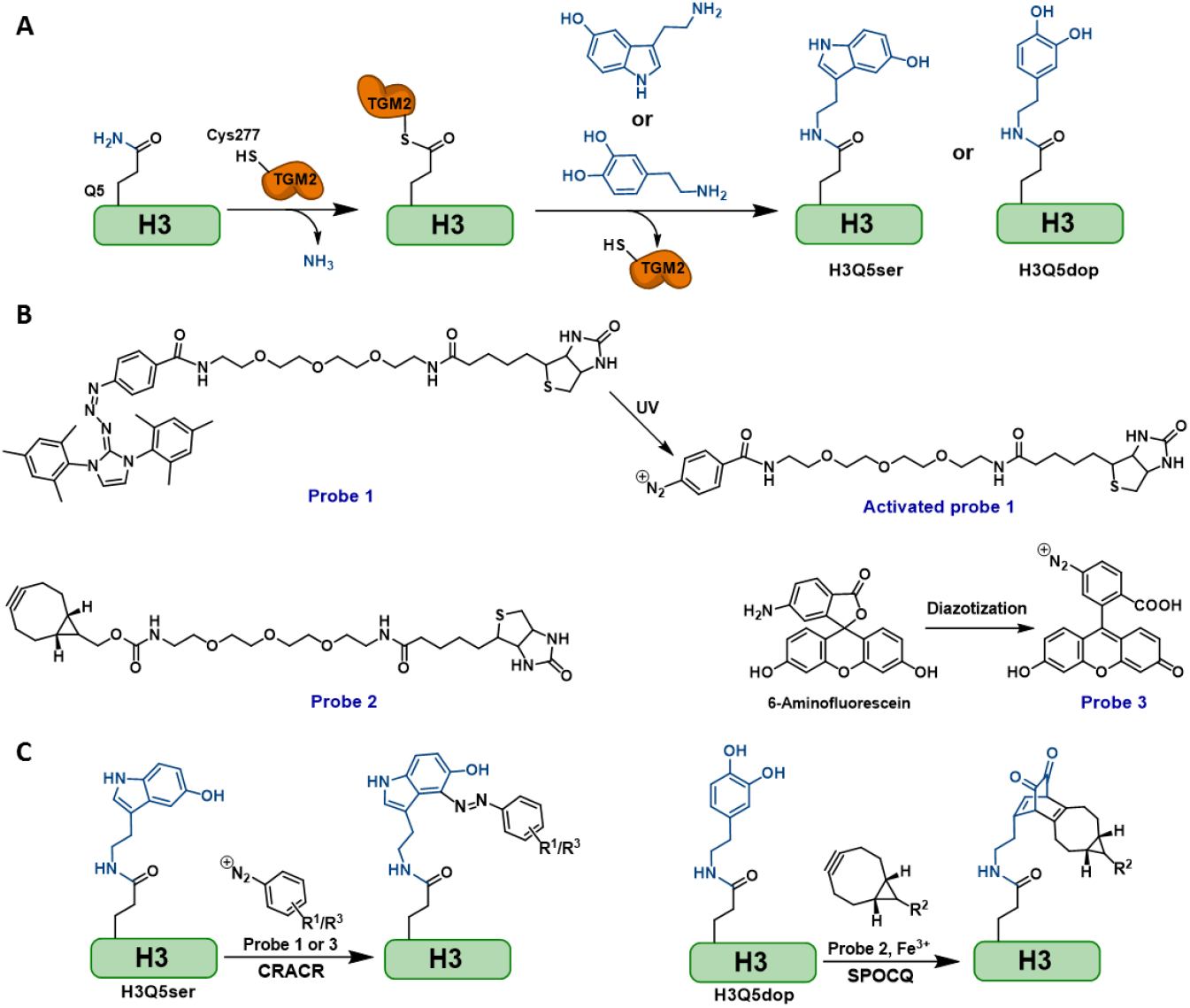
The strategy for bioorthogonal labeling and enrichment of histone monoaminylations in this study. (A) The biochemical mechanism of TGM2-catalyzed histone H3Q5 monoaminylation. (B) The structures of probes used in this study. (C) Bioorthogonal chemistry (CRACR and SPOCQ) used in this study for labeling and enrichment of monoaminylated histones.

To investigate the previously reported histone monoaminylation (*i*.*e*., H3Q5 serotonylation) in cultured cell samples, we treated the cells with 5-propargylated tryptamine (5-PT) and utilized Cu(I)-catalyzed azide-alkyne cycloaddition (CuAAC) to further label the introduced alkyne group with azido-fluorophore^17^ or biotin.^15^ However, the application of 5-PT required exogenous feeding to compete with endogenous serotonin in cells. Moreover, blocking the key functional hydroxyl group of serotonin^19^ may influence the binding of reader proteins to modified H3 tails, which is not an ideal mimic of histone serotonylation for downstream function studies. Due to these limitations, the development of a new generation of chemical probes is urgently needed to analyze endogenous histone monoaminylation levels in cells and tissues. One effective strategy is to design chemical biology probes that can specifically react with the modified residues of histones *via* bioorthogonal reactions. In this study, we designed and developed diazonium and bicyclononyne (BCN) probes to label and enrich histone serotonylation (H3Q5ser) and dopaminylation (H3Q5dop), respectively via chemoselective rapid azo-coupling reaction (CRACR)^20^ and strain-promoted oxidation-controlled cyclooctyne-1, 2-quinone cycloaddition (SPOCQ).^21^ Employing these powerful chemical tools, we have successfully uncovered that histone H3Q5 monoaminylations are enriched in breast and colorectal cancer tissues, which can exhibit regulatory functions through affecting H3K4 methylation and chromatin compaction status.

To specifically label and enrich histone serotonylation using the CRACR, we first synthesized a biotin-containing diazonium probe (Compound **1**; **Fig. 1B**) that was protected by a photoactive moiety, 1,3-dimesitylimidazolium, based on a reported synthetic route^20^ with slight modifications. The structure of this compound was thereafter confirmed by using nuclear magnetic resonance (NMR) and high-resolution mass spectrometry (HRMS) analyses (**Figs. S1** and **S2**). Probe **1** can release the diazonium group under ultraviolet (UV) treatment (λ=365 nm) to label C4 of serotonin in a site-specific and bioorthogonal manner (**Fig. 1C**). We first tested the reactivity of **1** using the monoaminylation-containing H3 peptides that were reported in our previous study^17^ and the liquid chromatography-mass spectrometry (LC-MS) analysis indicated that the H3Q5ser peptide was specifically modified by **1**, while the others did not react with **1** (**Figs. 2** and **S5**). Similarly, for the bioorthogonal labeling of histone dopaminylation, a BCN and biotin-containing probe (Compound **2**; **Fig. 1B**) was designed and synthesized, the structure of which was characterized by NMR and HRMS (**Figs. S3** and **S4**). *In vivo* biochemical assays with peptides as substrates and LC-MS analyses showed that the H3Q5dop peptide could be specifically derived by **2** *via* SPOCQ in the presence of Fe^3+^ (**Figs. 1C** and **2**). The bioorthogonally introduced biotin tags onto serotonylated and dopaminylated glutamine residues can be further utilized for imaging and pull-down enrichment.

**Figure 2.**
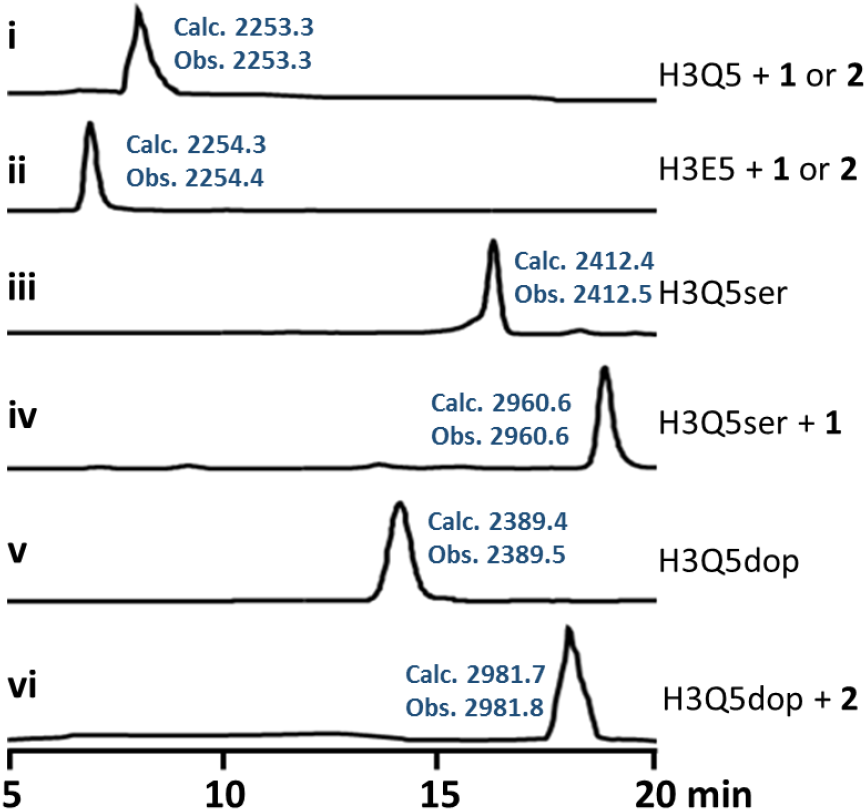
LC-MS analysis of modified and unmodified H3 peptides (λ = 214 nm): (i) H3Q5 (1-21) peptide treated by probe **1** or **2** (no reaction), (ii) H3Q5E (1-21) peptide treated by probe **1** or **2** (no reaction), (iii) H3Q5ser (1-21) peptide, (iv) H3Q5ser (1-21) peptide modified by probe **1**, (v) H3Q5dop (1-21) peptide, (vi) H3Q5dop (1-21) peptide modified by probe **2**. The synthesized H3 peptide sequence is NH_2_-ARTKQ(or E)TARKSTGGKAPRKQLA-COONH_2_.

To visualize the corresponding monoaminylations of H3Q5 on nucleosome core particles (NCPs), we prepared serotonylated and dopaminylated NCPs using recombinant TGM2 in the presence of Ca^2+^.^17^ The NCPs containing H3Q5E were used as negative controls that could not undergo TGM2-mediated monoaminylation. The probes **1** and **2** were incubated with serotonylated and dopaminylated wild-type NCPs, respectively, to introduce the biotin tags. Thereafter, the NCP histones were separated by using sodium dodecyl-sulfate polyacrylamide gel electrophoresis (SDS-PAGE) and imaged by the corresponding antibodies *via* immunoblotting. The signals of IRDye 680RD Streptavidin for biotin tags indicated that the two probes can successfully label and image H3Q5 monoaminylations in a bioorthogonal fashion (**Fig. 3A**). The site-specific antibodies for H3Q5ser and H3Q5dop are used as positive controls, which were produced in our previous studies.^15,16,17^ These customized antibodies worked well for the monoaminylated samples from *in vitro* assays, while they were not sensitive enough for cell or tissue samples from *in vivo* studies. Therefore, in our previous research, we utilized the 5-PT probe to pull down monoaminylated histones and anti-H3 antibody to indirectly detect the modification levels of H3.^15,17^ The results of *in vitro* experiments in this study showed that the bioorthogonal probes **1** and **2** are sensitive for labeling serotonylated and dopaminylated histones, which are competitive with the corresponding site-specific H3 monoaminylation antibodies (**Fig. 3B**). As the site-specific antibodies did not work efficiently for the direct visualization of histone monoaminylations in cultured cell lines, we applied the probes **1** and **2** to label the extracted histones from TGM2-overexpressed HEK 293T cells (in which the *tgm2* gene is silent) that had been treated with the corresponding monoamines.^17^ The signals of IRDye 680RD Streptavidin in the immunoblotting analyses indicated that **1** and **2** could precisely sense histone H3Q5 serotonylation and dopaminylation (**Fig. 3B**). In the meantime, 5-PT was utilized as a positive control to show the H3Q5 serotonylation catalyzed by wild-type TGM2 in cells.^15,17^ Notably, our data suggested that probe **1** could label not only serotonin, but 5-PT residues of modified proteins in a bioorthogonal fashion (**Fig. 3B**). In line with our previous *in vitro* and *in vivo* studies, the catalytically inactive mutation of TGM2, C277A, could not install the monoamines onto H3Q5 *via* transamination (**Fig. 3**).^17^

**Figure 3.**
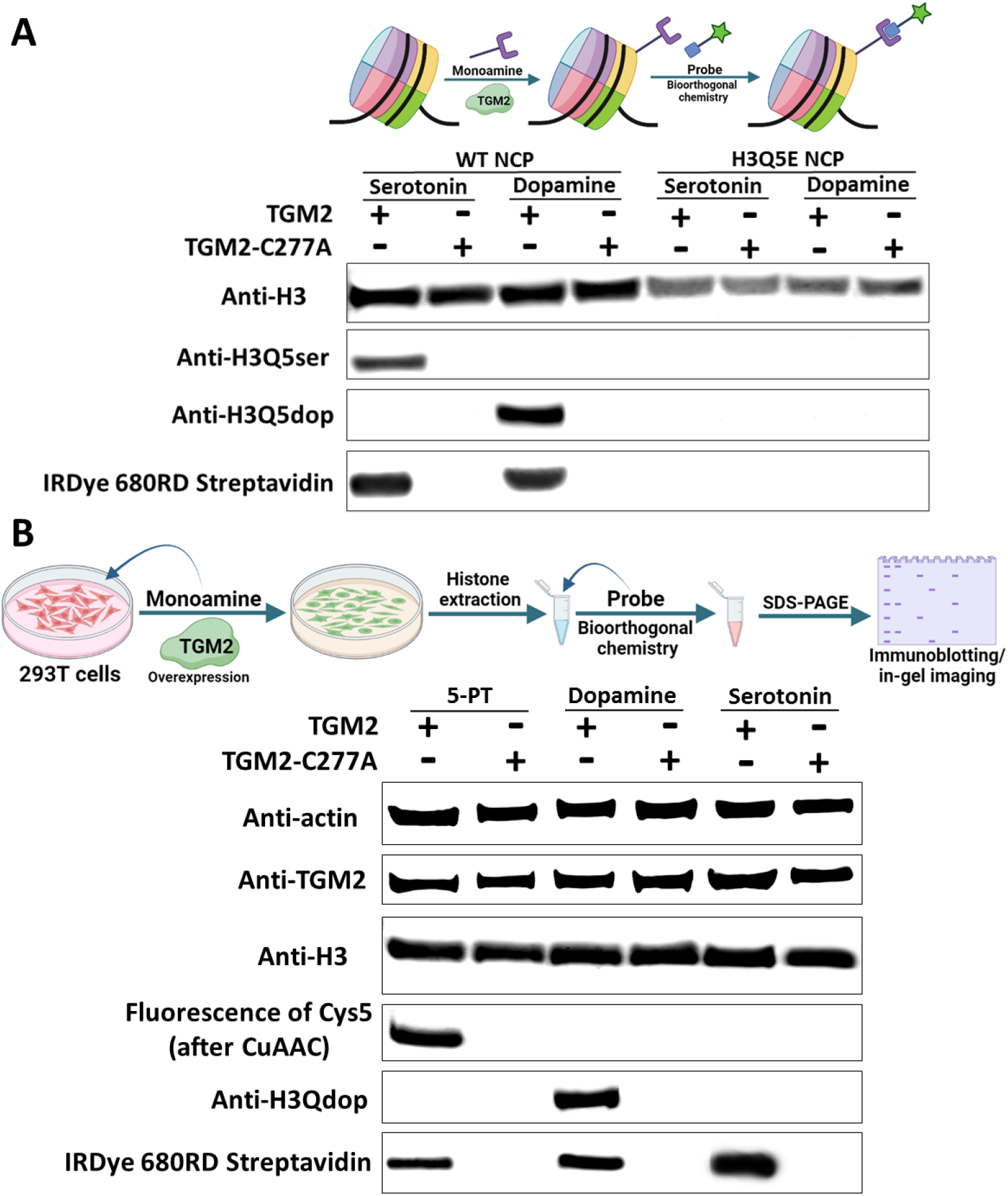
Immunoblotting analyses of monoaminylated histones from *in vitro* and *in cellulo* studies using probes **1** and **2**. (A) WT and H3Q5E NCPs were treated with TGM2-WT or TGM2-C277A in the presence of corresponding monoamine donors and then labeled by **1** or **2** to introduce the biotin tags. The NCP histones were separated by SDS-PAGE, followed by western blot analyses using the corresponding antibodies and IRDye 680RD Streptavidin (for indicating biotin tags). (B) TGM-2 and TGM2-C277A were overexpressed in HEK 293T cells in the the presence of corresponding monoamine donors (5-PT, dopamine, and serotonin). The histone fractions were extracted from the cells and separated by SDS-PAGE, followed by western blot analyses using the corresponding antibodies, Cy5 azide (for indicating the alkynyl group of 5-PT by CuAAC), and IRDye 680RD Streptavidin (for indicating biotin tags).

As TGM2 has been reported to be overexpressed in various types of cancers,^22^ we sought to apply the bioorthogonal probes to detect the histone monoaminylation levels in tumor tissues, the accumulation of which might be a consequence of TGM2 overexpression. To make probe **1** easier to use for the biomarker detection of tissue samples, we first transferred a commercial fluorophore, 6-aminofluorescein, to a fluorescent diazonium (Compound **3**) through a one-step diazotization reaction (**Fig. 1B**). Similar to probe **1, 3** can react with serotonylated histones in a bioorthogonal manner based on the same chemistry, CRACR.^20^ In comparison with **1, 3** is much easier to make and does not require UV-activation. Importantly, **3** can be directly used for fluorescent and in-gel imaging of serotonylated targets, which is convenient for tissue sample analyses and has potential for high-throughput screening assays. Given these tools, we extracted histones from breast cancer tissues from five patients^9,13^ and colorectal cancer tissues from C57BL/6J *Apc*^*Min/+*^ mice.^23^ Probe **3** was employed to visualize the H3Q5 serotonylation levels of these samples through in-gel imaging. The fluorescent signals indicated that histone serotonylation was highly accumulated in breast and colon tumors compared with the control non-tumor tissues (*i*.*e*., paracancerous tissues from the same five patients and colon tissues from the wild-type mice) (**Fig. 4**). The higher levels of histone serotonylation in these tumor tissues can be attributed to the overexpression of TGM2 in them (**Fig. 4**). Thereafter, immunoblotting analyses of probe **2**-treated histone fractions using IRDye 680RD Streptavidin further confirmed that histone dopaminylation was also unregulated in breast and colon tumors (**Fig. 4**).

**Figure 4.**
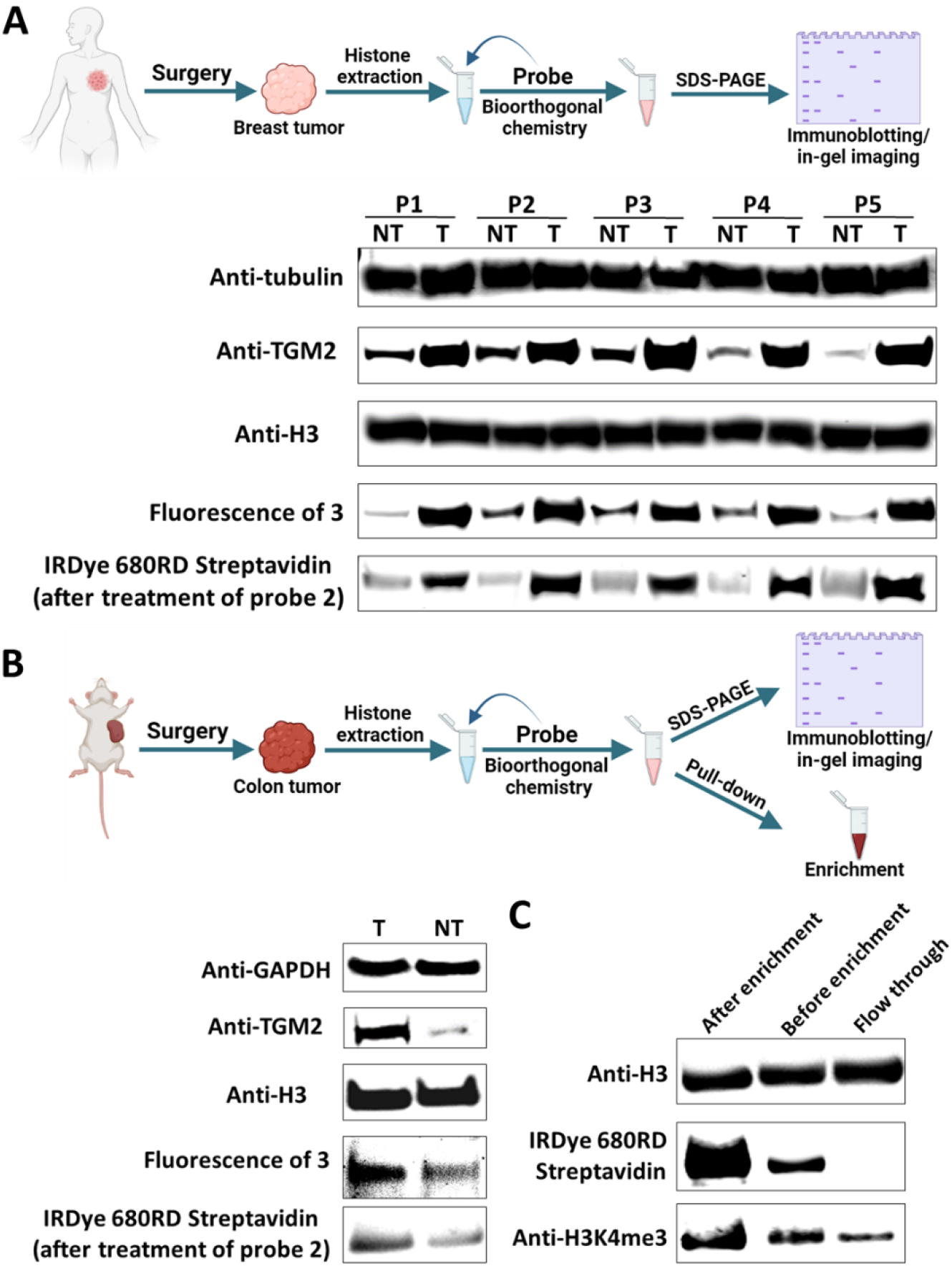
Immunoblotting and in-gel fluorescence imaging analyses of monoaminylated histones from *in vivo* studies using probes **2** and **3**. (A) The histone fractions were extracted from clinical tumor (T) and non-tumor (NT) samples from five different breast cancer patients (P1-P5) and treated with probes **2** and **3**. The histone fractions extracted from clinical tissues were separated by SDS-PAGE, followed by in-gel fluorescence (indicating the serotonylated H3 labeled by **3**) or western blot analyses using the corresponding antibodies. (B) The histone fractions were extracted from tumor (T) and non-tumor (NT) samples from the colon cancer mouse model and treated with probes **2** and **3**. The histone fractions extracted from mouse tissues were separated by SDS-PAGE, followed by in-gel fluorescence (indicating the serotonylated H3 labeled by **3**) or western blot analyses using the corresponding antibodies. (C) The histone fraction was extracted from tumor samples from the colon cancer mouse model and treated with probe **1**. Serotonylated histones were enriched by using streptavidin magnetic beads and analyzed by SDS-PAGE and western blot to uncover the crosstalk between H3Q5ser and H3K4me3.

Previous studies have shown that the metabolism of tryptophan to serotonin acts as a regulatory axis integrating the gut microbiome, diet, immunity, and colon cancer.^24,25^ Given the significant accumulation of histone serotonylation in colorectal cancer tissues, we sought to understand the functions of H3Q5 serotonylation in regulating the chromatin structures of cancer cells. In our studies, we have shown that H3Q5 serotonylation can potentiate binding to WDR5, which is a potential H3 “reader” protein and member of the MLL1 methyltransferase complex.^15,17,26^ The enhanced binding affinity between H3Q5ser and WDR5 can be attributed to multiple interactions, such as the cation-π stacking between R4828 of WDR5 and the indole ring of H3Q5ser.^17,27^ Although this enhanced binding had no effect on the enzymatic activity of MLL1 for H3K4 methylation, the existence of H3Q5ser could stabilize H3K4me3 from dynamic turnover by profoundly inhibiting the binding and activity of H3K4me3 erasers (such as KDM5B/C and LSD1).^26^ To uncover the crosstalk between H3Q5ser and H3K4me3 in colon tumor tissues, we incubated probe **1** and the extracted histone faction of colorectal cancer samples to introduce the biotin tag to serotonylated H3. Thereafter, probe **1**-modified H3Q5ser was enriched by streptavidin magnetic beads. H3K4me3 levels were then detected in three samples: probe **1**-enriched, unenriched, and flow-through histone factions from the mouse colon tumors. The subsequent immunoblotting analyses indicated that H3K4me3 levels were significantly higher in H3Q5ser-enriched samples that were pulled down by using probe **1** (**Fig. 4C**). These results suggested H3Q5 serotonylation could indeed stabilize H3K4me3 *in vivo*, which is a key the epigenetic mark in colon cancer development.^28,29^ Finally, the homododecameric (12-mer) nucleosomal arrays, which mimic the minimal chromatin fold,^9,13,30^ were applied to reveal the impacts of H3Q5ser on cellular three-dimensional chromatin architectures. Probe **3** was used to show that serotonin was installed to the wild-type nucleosomal arrays by TGM2 *in vitro* via in-gel imaging (**Fig. S6A**). Mg^2+^ precipitation analyses^9,13^ on the 12-mer arrays were performed to demonstrate that H3Q5 serotonylation decompacted the chromatin arrays (**Fig. S6B**), which could be attributed to the steric hindrance of H3Q5ser indole moiety. Overall, our results suggested H3Q5 serotonylation that was enriched in tumors could not only manipulate gene transcription via the crosstalk with H3K4me3, but also directly regulate the chromatin compaction states through the steric effect.

The biosynthesis of monoamines from indispensable amino acids (*e*.*g*., tryptophan to serotonin and phenylalanine to dopamine) is believed to act as a key hub connecting the gut microbiome, diet, signal transduction, and disease.^31,32^ As a newly identified epigenetic hallmark on histones, H3Q5 monoaminylation is an emerging link between amino acid metabolism and gene transcription.^15,16,17^ However, due to the lack of effective research tools (such as pan-specific antibodies and sensitive fluorophores for imaging), the pathophysiological relevance of H3Q5 monoaminylation still remains poorly understood. In this study, we designed and developed powerful chemical tools to label and enrich monoaminylated histones *via* bioorthogonal chemistry. The probes **1** and **2** can be applied for tracking and enriching histone monoaminylations, while **3** is a convenient and efficient tool for fluorescence imaging. Utilizing these tools, we have discovered that histone monoaminylation is highly enriched in breast and colon tumors, which plays a role in cancer epigenetic regulations through the crosstalk with H3K4 methylation (known as a key mark in cancer epigenetics)^33^ and steric hindrance.^34^ In summary, we observed for the first time that histone monoaminylations were accumulated in tumor tissues, which may serve as a biomarker for cancer screening and diagnosis in clinic. Moreover, we uncovered the essential epigenetic functions of H3Q5ser in cancer development, which included recruiting the “reader” WDR5,^27^ stabilizing the epigenetic mark H3K4me3,^26^ and directly decompacting the chromatin structures. Notably, the discoveries in this research further suggest that TGM2 may become a promising druggable target for cancer therapeutics.^35^

## Acknowledgements

This research work was financially supported by the NIH (R35 GM150676) and OSUCCC startup funds for Q.Z. The Zhu lab is supported by the NIH grant (R35 GM133510) for J.Z. The Maze lab is supported by the NIH grants (R01 DA056595 and R01 MH116900) and Howard Hughes Medical Institute for I.M.

