## Supplemental Information for "Bioorthogonal labeling and enrichment of histone monoaminylation reveal its accumulation and regulatory function in cancer cell chromatin"

[#] These authors contributed equally.

### MATERIALS AND METHODS

#### General methods (equipment, reagents, chemicals)

UV spectrometry was performed on a NanoDrop 2000c (Thermo Scientific). Biochemicals and media were purchased from Fisher Scientific or Sigma-Aldrich Corporation unless otherwise stated. T4 DNA ligase, DNA polymerase and restriction enzymes were obtained from New England BioLabs. PCR amplifications were performed on an Applied Biosystems Veriti Thermal Cycler using either Taq DNA polymerase (Vazyme Biotech) for routine genotype verification or Phanta Max Super-Fidelity DNA Polymerase (Vazyme Biotech) for high-fidelity amplification. Site-specific mutagenesis was performed according to standard procedures of the QuickChange Site-Directed Mutagenesis Kit purchased from Stratagene (GE Healthcare) or Mut Express II (Vazyme Biotech). Primer synthesis and DNA sequencing were performed by Integrated DNA Technologies and Genewiz, respectively. PCR amplifications were performed on a Bio-Rad T100TM Thermal Cycler. Centrifugal filtration units were purchased from Millipore, and MINI dialysis units purchased from Pierce. Size exclusion chromatography was performed on an AKTA FPLC system from GE Healthcare equipped with a P-920 pump and UPC-900 monitor. Sephacryl S-200 columns were obtained from GE Healthcare. All the western blots were performed using the primary antibodies annotated in **Supplementary Table 1** and fluorophore-labeled secondary antibodies annotated in **Supplementary Table 2** following protocols recommended by the manufacture. Blots were imaged on an Odyssey CLx Imaging System (Li-Cor). Amino acid derivatives and coupling reagents were purchased from AGTC Bioproducts. Dimethylformamide (DMF), dichloromethane (DCM) and triisopropylsilane (TIS) were purchased from Fisher Scientific and used without further purification. Hydroxybenzotriazole (HOBt) and O-(benzotriazol-1-yl)-N,N,N′,N′-tetramethyluronium hexafluorophosphate (HBTU) were purchased from Fisher Scientific. Trifluoroacetic acid (TFA) was purchased from Fisher Scientific. N,Ndiisopropylethylamine (DIPEA) was purchased from Fisher Scientific. Analytical reversed-phase HPLC (RP-HPLC) was performed on an Agilent 1200 series instrument with an Agilent C18 column (5 μm, 4 × 150 mm), employing 0.1% TFA in water (HPLC solvent A), and 90% acetonitrile, 0.1% TFA in water (HPLC solvent B) as the mobile phases. Analytical gradients were 0-70% HPLC buffer B over 45 minutes at a flow rate of 0.5 mL/minute, unless stated otherwise. Preparative scale purifications were conducted on an Agilent LC system. An Agilent C18 preparative column (15-20 μm, 20 × 250 mm) or a semi-preparative column (12 μm, 10 mm × 250 mm) was employed at a flow rate of 20 mL/min or 4 mL/min, respectively. HPLC Electrospray ionization MS (HPLC-ESI-MS) analysis was performed on an Agilent 6120 Quadrupole LC/MS spectrometer (Agilent Technologies). All immunoblotting experiments in this research were performed at least 3X. For the synthesis of probe molecules used in this study, all commercial chemicals were purchased from Sigma Aldrich, TCI chemicals, AK Scientific, Fischer Scientific, Broadpharm and used without further purification. The reagents and solvents were handled following the safety processes required as instructed by the manufacturer. Organic and aqueous waste was disposed of following standard safety protocols. Solvents for workup were purchased from Fisher Chemical, and anhydrous solvents were purchased from Sigma Millipore in a sealed bottle and degassed by passing $N_2$ before each use. Reaction progress was monitored by using normal phase TLC silica gel 60 $F_{254}$ plates by Sigma Aldrich (aluminum backed 20 X 20 cm). Developed plates were analyzed by visualizing under a UV-light and/or staining with Phosphomolybdic Acid (PMA) Stain (100 mL absolute ethanol and 10 g PMA). Isolation and purification of the crude reaction materials were performed using silica gel ($SiO_2$) by Acros Organic (0.030-0.200 mm, 60 $A^0$). Organic solvents were removed under vacuum using a Heidolph Rotavapor equipped with a dry ice condenser. Deuterated solvents (such as $CDCl_3$) for NMR characterization were purchased from Sigma Aldrich. $^1H$ NMR, $^{13}C$ NMR data were recorded on a Bruker Avance-600 MHz spectrometer at 22 °C. Chemical shifts (δ) are reported in ppm to the internal standard of residual $CDCl_3$ (δ7.26: $^1H$ NMR, δ 77.16: $^{13}C$ NMR). All the $^1H$ NMR are reported as follows; chemical shift (δ ppm), multiplicity (s, singlet; d, doublet; t, triplet; q, quartet; m, multiplet; dd, doublet of doublets), coupling constant (Hz), integration and assigned proton. Data for $^{13}C$ NMR spectroscopy are reported in chemical shift (δ ppm). High-resolution mass spectrometry (HRMS) analyses of the molecules were performed on a Thermofisher Scientific Q Exactive (QE) LC-MS/MS.

**Synthesis of probe 1**

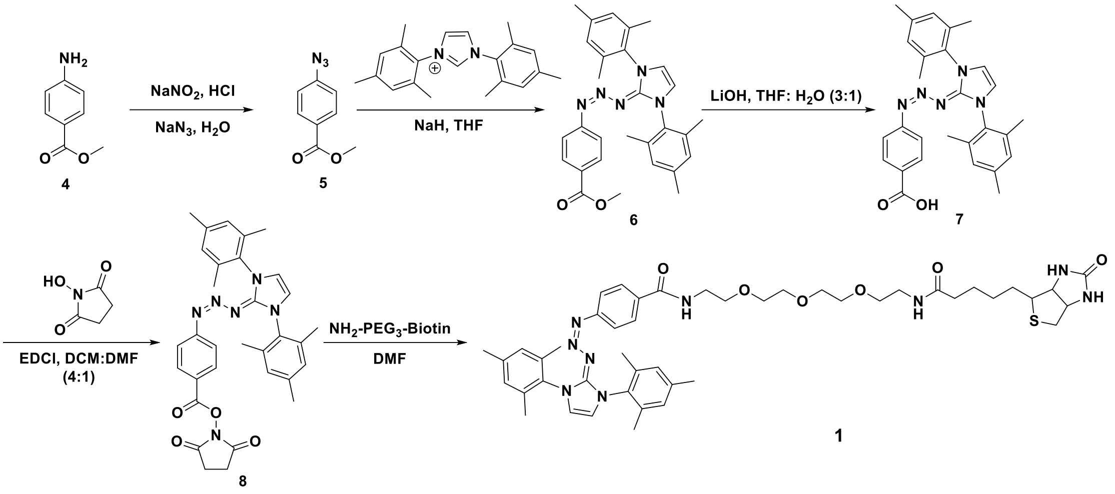


The probe **1** was synthesized following a reported syntactic route with slight modifications.[1] The modified synthesis started with commercially available methyl-4-amino benzoate (**4**), where the amino group was converted to the azido group in the presence of acidic solutions of sodium nitrite ($NaNO_2$) and sodium azide ($NaN_3$). Thereafter, a trizabutadiene group was installed to methyl 4-azidobenzoate (**5**) utilizing 1,3-dimesitylimidazolium chloride and sodium hydride (NaH). The ester group of **6** was then deprotected with aqueous lithium hydroxide (LiOH) to generate compound **7**. The carboxylic acid group of **7** was subsequently activated with *N*-hydroxysuccinamide and 1-(3-Dimethylaminopropyl)-3-ethylcarbodiimide hydrochloride hydrochloride (EDCl) to give **8**. At last, compound **8** was conjugated with biotin-$PEG_3$-amine to get the photoactive probe **1**. Compound **1** was purified through recrystallization for the NMR analysis and described applications in this study.

$^1H$ NMR (600 MHz, $CDCl_3$) δ 7.46 (d, *J* = 8.3 Hz, 2H), 7.07 (broad, 1H, NH), 6.95 (s, 4H), 6.79 (broad, 1H, NH), 6.60 (s, 2H), 6.52 (d, *J* = 8.3 Hz, 2H), 6.48 (broad, 1H, NH), 5.59 (broad, 1H, NH), 4.33 (dd, *J* = 8.0, 4.9 Hz, 1H), 4.13 (dd, *J* = 8.2, 4.7 Hz, 1H), 3.62 – 3.48 (m, 13H), 3.43 (t, *J* = 5.3 Hz, 2H), 3.32 – 3.27 (m, 2H), 2.99 (td, *J* = 7.3, 4.5 Hz, 1H), 2.75 (dd, *J* = 12.8, 4.9 Hz, 1H), 2.62 (s, 1H), 2.32 (s, 6H), 2.10 – 2.07 (m, 14H), 1.66 – 1.60 (m, 1H), 1.57–1.50 (m, 3H), 1.31 (p, *J* = 7.6 Hz, 2H).

$^{13}C$ NMR (150 MHz, $CDCl_3$) δ 173.5, 171.2, 167.5, 164.0, 139.1, 134.9, 133.7, 130.9, 129.4, 128.1, 127.4, 120.7, 117.5, 70.4, 70.3, 70.2, 70.0, 69.9, 69.9, 61.7, 60.4, 60.2, 55.6, 40.5, 39.7, 39.0, 35.9, 28.1, 21.1, 17.9, 14.2.
HRMS (ESI-TOF-MS) (m/z): $[M+H]^+$ Calculated for $C_{46}H_{62}N_9O_6S$ 868.4544 obtained 868.4495.

**Synthesis of probe 2**

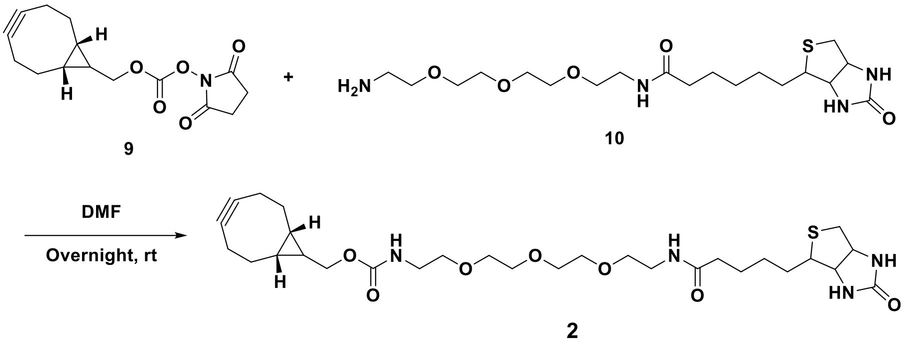


Probe **2** is a newly reported compound in this study. The starting material, compound **9** (100 mg, 0.3432 mmol, 1 eq), was dissolved in DMF (3 mL). Thereafter, biotin-PEG$_3$-amine, **4** (172.44 mg, 0.41 mmol, 1.2 eq), was added into the solution. The reaction mixture was washed with water (3 X 10 mL) and extracted with DCM (3 X 10 mL) after stirring overnight at room temperature. The organic layers were dried over $Na_2SO_4$ and then concentrated under reduced pressure. Preparative RP-HPLC was used to purify probe **2** for the NMR analysis and described applications in this study. Purified compound **2** turned to be transparent oil with a yield of 55% and $R_f$ 0.4 in 1:10 MeOH-DCM.
$^1H$ NMR (600 MHz, Chloroform-d) δ 6.80 (s, 1H), 6.57 (s, 1H), 5.72 (s, 1H), 5.40 (s, 1H), 4.48-4.46 (q, *J*= 6 Hz, 1H), 4.29-4.27 (q, *J*= 6 Hz, 1H), 4.12-4.10 (d, *J*= 12 Hz, 1H), 3.60 (s, 8H), 3.55-3.52 (q, *J*= 6 Hz, 4H), 3.41-3.39 (m, 2H), 3.34 (s, 2H), 3.13-3.09 (q, *J*= 12 Hz, 1H), 2.88-2.85 (dd, *J*= 12, 6 Hz, 1H), 2.26-2.17 (m, 8H), 1.74-1.68 (m, 1H), 1.67-1.61 (m, 3H), 1.59-1.54 (m, 2H), 1.43-1.38 (m, 2H), 1.35-1.30 (m, 1H), 0.92-0.89 (t, *J*= 6 Hz, 2H)
$^{13}C$ NMR (150 MHz, $CDCl_3$) *δ* 98.9, 70.4, 70.2, 70.1, 62.8, 61.9, 60.3, 55.7, 40.6, 39.2, 29.1, 28.3, 28.1, 25.7, 21.5, 20.2, 17.8
HRMS (ESI-TOF-MS) (m/z): $[M+H]^+$ Calculated for $C_{29}H_{47}N_4O_7S$ 595.3165 obtained 595.3165.

**Synthesis of probe 3**

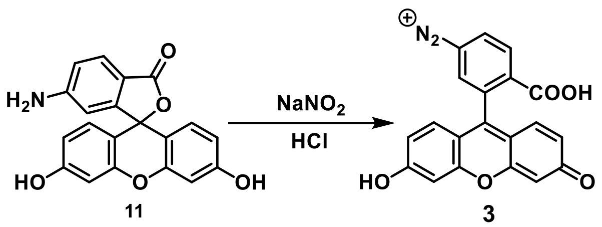


Probe **3** was synthesized from a commercial fluorophore, 6-aminofluorescein (**11**), through a one-step diazotization reaction in aqueous solution, which needed to be

freshly made before using due to its instability.[1] Briefly, 20 µL of $NaNO_2$ (60 mM) (Sigma) was added to 100 µL of 6-aminofluorescein (10 mM) (Sigma), which was dissolved in 10mM HCl. After a quickly vortex, the mix was kept on ice to generate the solution of fluorescein diazonium **3** (8.3 mM). At last, the solution was diluted to 1.5 mM with water for the further applications in this study.
HRMS (ESI-TOF-MS) (m/z): $M^+$ Calculated for $C_{20}H_{11}N_2O_5^+$ 359.0662 obtained 359.0663.

**Recombinant histone expression and purification**
Recombinant human histones H2A, H2B, H3 (wild-type and H3Q5E mutant) and H4 were expressed in *E. coli* BL21 (DE3) or *E. coli* C41 (DE3), extracted by guanidine hydrochloride and purified by flash reverse chromatography, as previously described.[2] The purified histones were analyzed by RP-LC-ESI-MS.[2]

**Preparation of histone octamers and '601' DNA**
Octamers were prepared as previously described.[2] Briefly, recombinant histones were dissolved in unfolding buffer (20 mM Tris-HCl, 6M GdmCl, 0.5mM DTT, pH 7.5), and combined with the following stoichiometry: 1.1 eq. H2A, 1.1 eq. H2B, 1 eq. H3.2, 1 eq. H4. The combined histone solution was adjusted to 1 mg/mL concentration and transferred to a dialysis cassette with a 7000 Da molecular cutoff. Octamers were assembled by dialysis at 4 ℃ against 3 × 1 L of octamer refolding buffer (10 mM Tris-HCl, 2 M NaCl, 0.5mM EDTA, 1 mM DTT, pH 7.5) and subsequently purified by size exclusion chromatography on a Superdex S200 10/300 column. Fractions containing octamers were combined, concentrated, diluted with glycerol to a final 50% v/v and stored at -20 ℃. The 147-bp 601 DNA fragment was prepared by digestion from a plasmid containing 30 copies of the desired sequence (flanked by blunt EcoRV sites on either site) and purified by PEG-6000 precipitation as described before.[2]

**Mononucleosome assembly**
The mononucleosome assembly was performed according to the previously described salt dilution method with slight modification.[2] Briefly, the purified wild-type octamers were mixed together with 601 DNA (1:1 ratio) in a 2 M salt solution (10 mM Tris pH 7.5, 2 M NaCl, 1 mM EDTA, 1 mM DTT). After incubation at 37 ℃ for 15 min, the mixture was gradually diluted (9 × 15 min) at 30 ℃ by dilution buffer (10 mM Tris pH 7.5, 10 mM NaCl, 1 mM EDTA, 1 mM DTT). The assembled mononucleosomes were concentrated and characterized by native gel electrophoresis (5% acrylamide gel, 0.5 × TBE, 120 V, 40 minutes) using ethidium bromide (EtBr) staining.

**Nucleosomal array assembly**
Dodecameric repeats of the 601 DNA sequence separated by 30-bp linkers were produced from pWM530 using EcoRV digestion and PEG-6000 precipitation according to the published procedure.[3] Homotypic dodecameric arrays were assembled from purified octamers and recombinant DNA in the presence of buffer DNA (MMTV) by salt gradient dialysis as previously described. The resulting arrays were purified and concentrated using $Mg^{2+}$ precipitation at 4 °C.[3]

**Expression of recombinant wild-type TGM2 and C277A mutant**
The wild-type and mutant TGM2 were expressed based on our previous protocols.[4] Briefly, the $His_8$-tagged proteins were expressed in *E. coli* Rosetta (DE3) cells with an overnight IPTG induction at 16 ℃. The bacterial pellet was lysed by sonication and the

lysate was cleared by centrifugation at 12,000 r.p.m. for 30 minutes. The lysate was loaded on HisTrap HP Column (GE Healthcare) and eluted on the AKTA FPLC, followed by desalting using Zeba Spin Desalting Columns (7 K MWCO, 10 mL) according to the manufacturer's protocol. Purified recombinant proteins were analyzed by SDS-PAGE and concentrated using stirred ultrafiltration cells (Millipore) according to the manufacturer's protocol. The concentration of each protein was determined using 280 nm wavelength on a NanoDrop 2000c (Thermo Scientific).

**Peptide synthesis**

Standard Fmoc-based Solid Phase Peptide Synthesis (FmocSPPS) was used for the synthesis of peptides in this study. Generally, the peptides were synthesized on ChemMatrix resins with Rink Amide to generate C-terminal amides. Peptides were synthesized using manual addition of the reagents (using a stream of dry $N_2$ to agitate the reaction mixture). For amino acid coupling, 5 eq. Fmoc protected amino acid were pre-activated with 4.9 eq. HBTU, 5 eq. HOBt and 10 eq. DIPEA in DMF and then reacted with the N-terminally deprotected peptidyl resin. Fmoc deprotection was performed in an excess of 20% (v/v) piperidine in DMF, and the deprotected peptidyl resin was washed thoroughly with DMF to remove trace piperidine. Cleavage from the resin and side-chain deprotection were performed with 95 % TFA, 2.5% TIS and 2.5% $H_2O$ at room temperature for 1.5 hours. The peptides were then precipitated with cold diethyl ether, isolated by centrifugation and dissolved in water with 0.1 % TFA followed by RP-HPLC and ESI-MS analyses. Preparative RP-HPLC was used to purify the peptides of interest.[2,3]

For the synthesis of site-specific monoaminylated H3 peptides, Fmoc-Glu(OAll)-OH was incorporated at position 5 for the orthogonal deprotection and further monoaminylation. Briefly, the peptides were deprotected by $Pd(PPh_3)_4$ and $PhSiH_3$ on resins and then conjugated with monoamine donors (*i.e.*, serotonin hydrochloride, and acetonide-protected dopamine) via PyAOP and DIEA catalysis.[4]

***In vitro* labeling assays**

For H3 peptide labeling assays, 2 mM peptides were generally treated with 10 mM probe molecules on ice (probe **1**) or at room temperature (probe **2**) for 5 min and then analyzed by LC-MS. Specifically, purified compound **1** was dissolved in DMSO and was then photolyzed for 1 min using a 43 W UV light (Kessil PR160L-370nm), in which time the conversion of the triazabutadiene to diazonium was confirmed to be complete. Peptides were dissolved in 100 mM phosphate buffer (pH 7) and incubated with freshly photolyzed **1**, followed by the addition of 5-HTP (10mM) to quench the unreacted diazonium; Compound **2** was dissolved in DMSO and then added to the peptides that had been pre-treated with 4 mM $K_3Fe(CN)_6$ for 5min in 1 X PBS (pH 8). After 5 min at room temperature, the reaction was quenched by adding 10 mM dopamine and the resulting product was analyzed by LC-MS.

For nucleosome core particle (NCP) labeling assays, 1 μM NCPs were treated with 0.1 μM TGM2 in the buffer (pH 7.5) containing 50 mM Tris-HCl, 5 mM $CaCl_2$, and 2 mM DTT (freshly added) at 37 ℃ in the presence of corresponding monoamaines (0.5 μM) for 2 hours. The buffer exchange for the further labeling assays was performed using 0.5 mL Centrifugal Filter (3K, Millipore) with a 120-fold v/v for the removal of excess monoamine from the old reaction buffer systems. Specifically, the modified NCPs were dissolved in 100 mM phosphate buffer (pH 7) via buffer exchange and then incubated with freshly photolyzed **1** (10 μM) on ice for 10 min, followed by the addition of 5-HTP (10 μM) to quench the unreacted diazonium; Compound **2** was added to the modified NCPs that had been pre-treated with 4 μM $K_3Fe(CN)_6$ for 10 min in 1 X PBS (pH 8). After 10 min at room temperature, the reaction was quenched by adding 10 μM dopamine. The labeled NCPs were analyzed by sodium dodecyl sulfate polyacrylamide gel electrophoresis (SDS-PAGE) followed by western blot analysis using IRDye 680RD Streptavidin (LI-COR). H3 was used as the loading control in SDS-PAGE and western blot analyses.

For the labeling of modified 12-mer nucleosomal arrays, 0.1 μM arrays were treated with 0.1 μM TGM2 in the buffer (pH 7.5) containing 50 mM Tris-HCl, 5 mM CaCl2, and 2 mM DTT (freshly added) at 37 ℃ in the presence of corresponding monoamaines (0.5 μM) for 2 hours. The modified arrays were then dissolved in 100 mM phosphate buffer (pH 7) via buffer exchange and incubated with freshly prepared probe **3** (20 μM) on ice for 10 min, followed by the addition of 5-HTP (20 μM) to quench the unreacted diazonium. The labeled arrays were analyzed by agarose-polyacrylamide gel electrophoresis (APAGE) followed by in-gel fluorescence imaging. EtBr was used to stain the DNA of arrays as the loading control in APAGE analyses.

For the Cy5 labeling of 5-PT-modified histones from cultured cells, extracted histones were desalinated, lyophilized and then resuspended in DPBS buffer containing 0.4 % SDS. 50 μL of freshly dissolved histones was added to a premixed solution containing 3 μL of 10 mM Cy5-azide (Sigma-Aldrich, 777323), 10 μL of a 3:7 mixture of 50 mM $CuSO_4$ and 100 mM THPTA, and then vortexed. Thereafter, 5 μL of 100 mM freshly made TCEP was added to initiate the click reaction followed by incubation (1-2 hours) at 30 $^{o}$C. Then, 10 μL of 0.5 M EDTA was added to quench the reactions. Excess reagents were removed by $MeOH/CHCl_3$ protein precipitation or concentration-dilution using a 0.5 mL Centrifugal Filter (3K, Millipore). Pellets were washed by 500 μL MeOH/H2O (9:1) prior to a second centrifugation. The air-dried protein samples were then analyzed by SDS-PAGE followed by in-gel imaging using Odyssey CLx Imaging System (wavelength 680 nm).[4] Similarly, for the biotin labeling of serotonin- or dopamine-modified histones from cultured cells using probes **1** and **2**, 0.5 mM probes were respectively used to incubate with histones dissolved in 100 mM phosphate buffer (pH 7) and 1 X PBS (pH 8) on ice and room temperature for 20 min before the reactions were quenched by 5-HTP (0.5 mM) and dopamine (0.5 mM), followed by SDS-PAGE and western blot analyses.

**Expression of TGM2 in HEK 293T cells**

The plasmid of TGM2 and its mutant were constructed expressed in HEK 293T cells based on the protocol of our previous research.[4] Briefly, wild-type TGM2 and the TGM2-C277A mutant were overexpressed in HEK 293T cells using Lipofectamine 2000 Transfection Reagent (Thermo Fisher Scientific) according to the manufacturer's protocol. HEK 293T cells (ATCC) were cultured at 37 ℃ with 5% $CO_2$ in DMEM medium supplemented with 10% fetal bovine serum (FBS) (Sigma-Aldrich), 2 mM L-glutamine and 500 units $mL^{-1}$ penicillin and streptomycin. The cells were stimulated with 2 μM calcium ionophore (Sigma-Aldrich, A23187) for 6 h at 37 ℃ before lysis in DPBS buffer (Gibco), and then the expression of TGM2 was detected by western blot analyses with anti-TGM2 antibody (CST, #3557).

**Extraction of histones from cultured cells**

The extraction of histones from cells was performed according to the previously described high salt extraction method.[2] Briefly, the cell lysis solution was prepared using extraction buffer (10 mM HEPES pH 7.9, 10 mM KCl, 1.5 mM $MgCl_2$, 0.34 M sucrose, 10% glycerol, 0.2% NP40, protease and phosphatase inhibitors to 1 × from stock). After spinning down, the pellet was extracted using a no-salt buffer (3 mM EDTA, 0.2 mM EGTA). After discarding the supernatant, the final pellet was extracted by using high-salt buffer (50 mM Tris pH 8.0, 2.5 M NaCl, 0.05% NP40) in 4 ℃ cold room for 1 hour. After spinning down, the supernatant containing extracted histones was collected, desalted, and lyophilize for further labeling assays and analyses.[4]

**Cell fractionation**

Cytosolic and nuclear fractions were prepared using NEPER Nuclear and Cytoplasmic Extraction Reagents (Thermo Scientific) according to the manufacturer's protocol. Histones were extracted from the pellet using high salt extraction protocol, as described above (1). Purity of fractionation was evaluated using the following antibodies: anti-Actin (cytosol), anti-MEK ½ (nucleoplasm) and anti-H3 (chromatin).[2,3]

**Protein extraction from tumor tissues**

The breast tumor tissues from patients were reported in our previous studies and processed using the same protocol.[2,3] For obtaining the colon tumor tissues, breeding pairs of wild-type (WT) and C57BL/6J $Apc^{Min/+}$ mice were purchased from the Jackson laboratory. Six-week-old WT female mice was breed with six-week-old male $Apc^{Min/+}$ mice. All pups were genotyped following the protocol on the Jackson Laboratory. All studies were performed in compliance with institutional guidelines under an Institutional Animal Care and Use Committee-approved protocol (2018A00000089). Colons and intestines' tumor from all $Apc^{Min/+}$ mice at age 20 weeks were collected on ice and isolated by surgical blades.[5]

The cytoplasmic and nuclear proteins from the mouse colon tissues were extracted by using Cytoplasmic and Nuclear Protein Extraction Kit (BOSTER BIO) according to the manufacturer's protocol. The mixture of cytoplasmic and nuclear protein fraction was used for immunoblotting analysis of TGM2 expression, and the pellet left was used for histone extraction. The extraction of histone was performed according to the previously described acid extraction method.[2,3,4] Briefly, the chromatin pellet was resuspended in 0.4 N $H_2SO_4$ and rotated at 4 $^oC$ overnight to extract histones. After centrifugation at 16,000 X g for 10 minutes, the supernatant was transferred to a new tube followed by adding 132 μL of 100% TCA to precipitate histones overnight. At last, histones were dried at room temperature after precipitation by centrifugation and being washed with 1 mL cold acetone. The dried histones were then dissolved in 100 μL Milli-Q water. The concentration of each sample was determined using 280 nm wavelength on a Quickdrop (MOLECULAR DEVICES). The histone solution was then diluted to ~0.5 μg/μL for the further labeling assays and analyses.

**Labeling of modified histones from tumor tissues**

Freshly prepared probe **3** solution was used for the labeling experiments of serotonin-modified histones from tumor tissues. Briefly, 150 μM probe **3** was added to 100 mM phosphate buffer (pH 7) containing 40 μg histones and incubated on ice for 15 min. Unreacted diazonium was then quenched by adding 150 μM 5-HTP. At last, the labeled histones were analyzed using SDS-PAGE, western blot, and in-gel fluorescence imaging. The in-gel imaging was conducted by using UV wavelength on GelDoc Go Imaging System (BIO-RAD).

For the labeling assays using probe **2**, 70 μL of 35 μg histones were dissolved in 10 μL 10 X PBS (pH 8) and then incubated with 10 μL of $K_3Fe(CN)_6$ (0.1 mM) at room temperature for 10 minutes followed by adding 10 μL of probe 2 that was dissolved in DMSO (5mM). After 30 minutes of reaction at room temperature, the reaction was quenched by adding 10 μL of 5 mM dopamine. At last, the labeled histones were analyzed using SDS-PAGE and western blot. Western blot was performed according to the previously protocol.[2,3,4] Briefly, proteins were analyzed using 12% SDS-PAGE gels, which were transferred from the gel to a PVDF membrane (Life Technologies) using a membrane transfer system (Bio-Rad). The membrane was blocked with 5% milk for 1 h at room temperature with gentle swinging and incubated with H3 first antibody at 4 $^{o}$C overnight after 4 time washing (5 minutes per time). On the next day, antibody solution was removed, and membrane was incubated with a mixture of IRDye 680RD Streptavidin (1:15000; LI-COR) and the second antibody (1:10000) at room temperature for 1 hour. The membrane was washed 4 times before signal detection by the imaging system (ODYSSEY CLX).

**Statistics and reproducibility**

All *in vitro*, *in cellulo*, and *in vivo* western blotting and mass spectrometry analyses were repeated independently at least three times with similar results. Significance was determined at $p<0.05$. All data are represented as mean ± SEM. Statistical analyses were performed in GraphPad Prism 9. The convolution of LC-MS data in this study was performed using SAMMI and MassWorks (Cerno Bioscience).

### SUPPLEMENTARY TABLES

| Host | Epitope | WB | Vendor |
|---|---|---|---|
| **Rabbit** | Anti-TGM2 | 1:500 (tumor) 1: 1000 (*in vitro* and *in cellulo*) | CST (3557S) |
| **Chicken** | Anti-H3 | 1: 1000 | Abcam (ab134198) |
| **Mouse** | Anti-H3 | 1: 1000 | Abcam (ab10799) |
| **Mouse** | Anti-Actin | 1: 1000 | CST (3700S) |
| **Rabbit** | Anti-H3Q5ser | 1: 1000 | Millipore (ABE1791) |
| **Rabbit** | Anti-H3Q5dop | 1: 1000 | Millipore (ABE2588) |
| **Rabbit** | Anti-H3K4me3 | 1:1000 | Abcam (ab8580) |
| **Rabbit** | Anti-H3 | 1:50000 (tumor) | Abcam (ab1791) |

**Supplementary Table 1.** Primary antibodies used in this study.

| Host | Epitope | Label | Dilution | Vendor |
| --- | --- | --- | --- | --- |
| Donkey | Anti-Chicken | IRDye 800CW | 1: 15000 | Li-Cor |
| Goat | Anti-Mouse | IRDye 680RD | 1: 15000 | Li-Cor |
| Goat | Anti-Mouse | IRDye 800CW | 1: 15000 | Li-Cor |
| Goat | Anti-Rabbit | IRDye 800CW | 1: 15000 | Li-Cor |
| Goat | Anti-Rabbit | IRDye 680RD | 1: 15000 | Li-Cor |

**Supplementary Table 2.** Secondary antibodies used in this study.

### SUPPLEMENTARY FIGURES AND LEGENDS

**Figure S1.** $^{1}$H (top) and $^{13}$C (bottom) NMR spectrum of compound **1**.

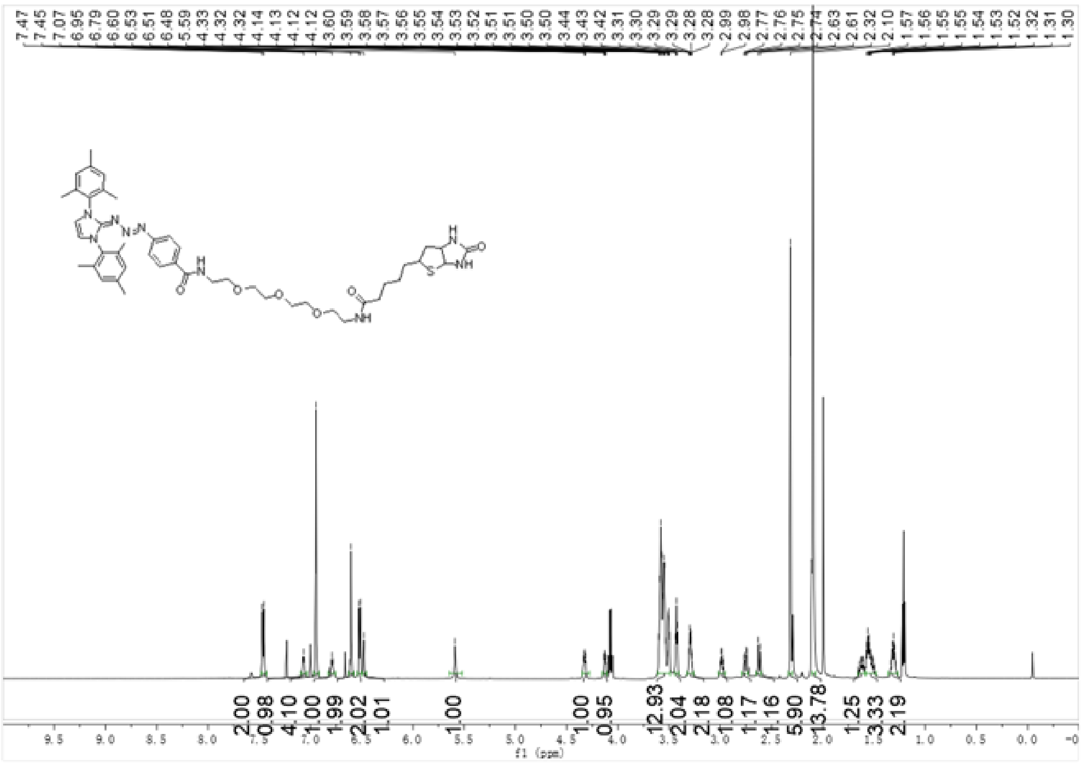


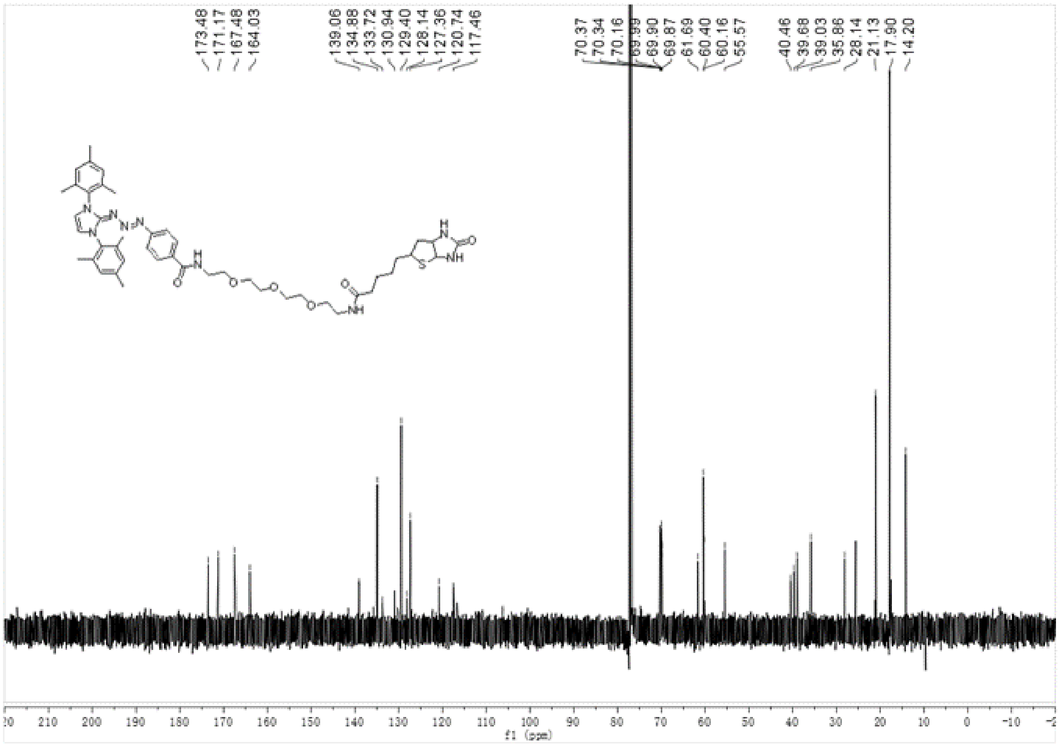

**Figure S2.** HRMS spectrum of compound **1**.

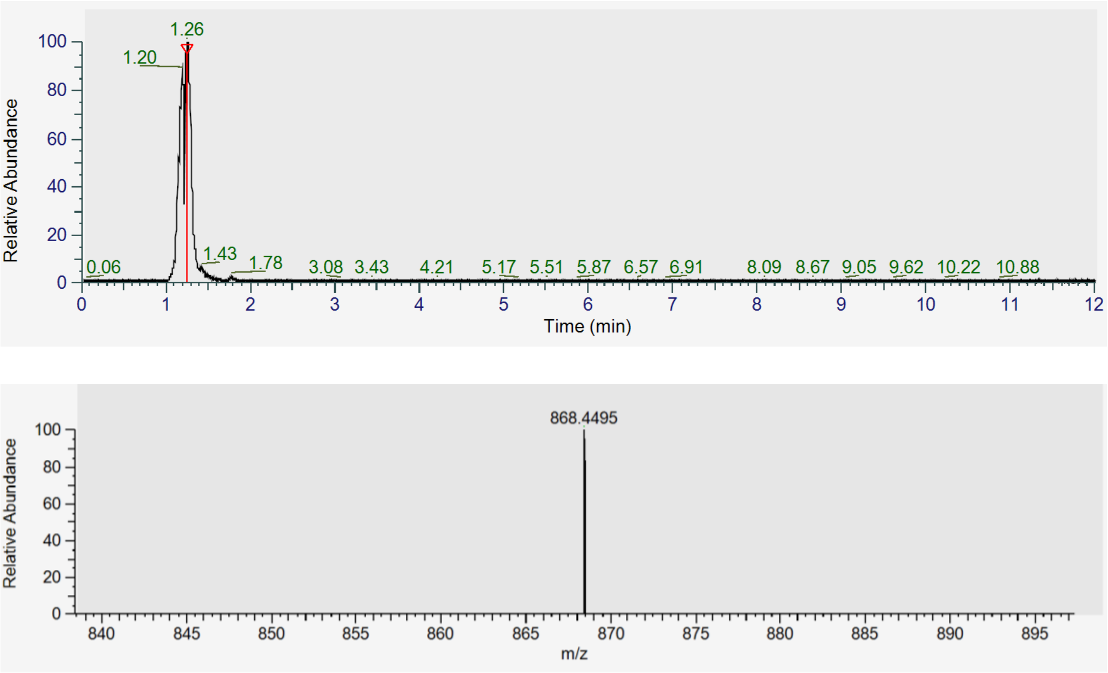

**Figure S3.** $^{1}$H (top) and $^{13}$C (bottom) NMR spectrum of compound **2**.

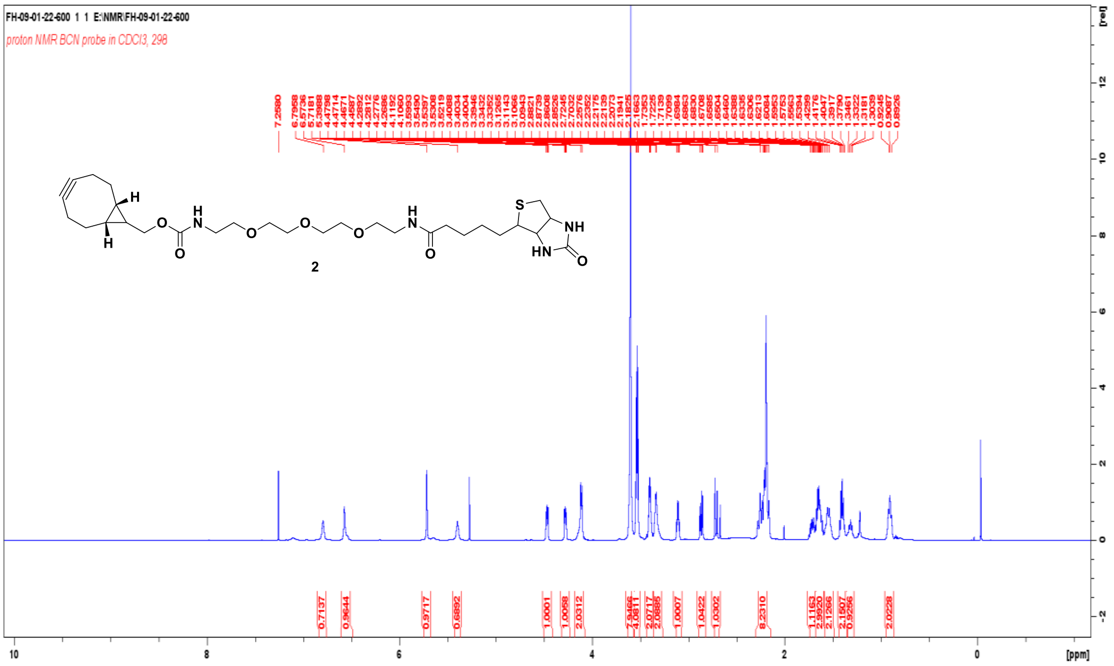


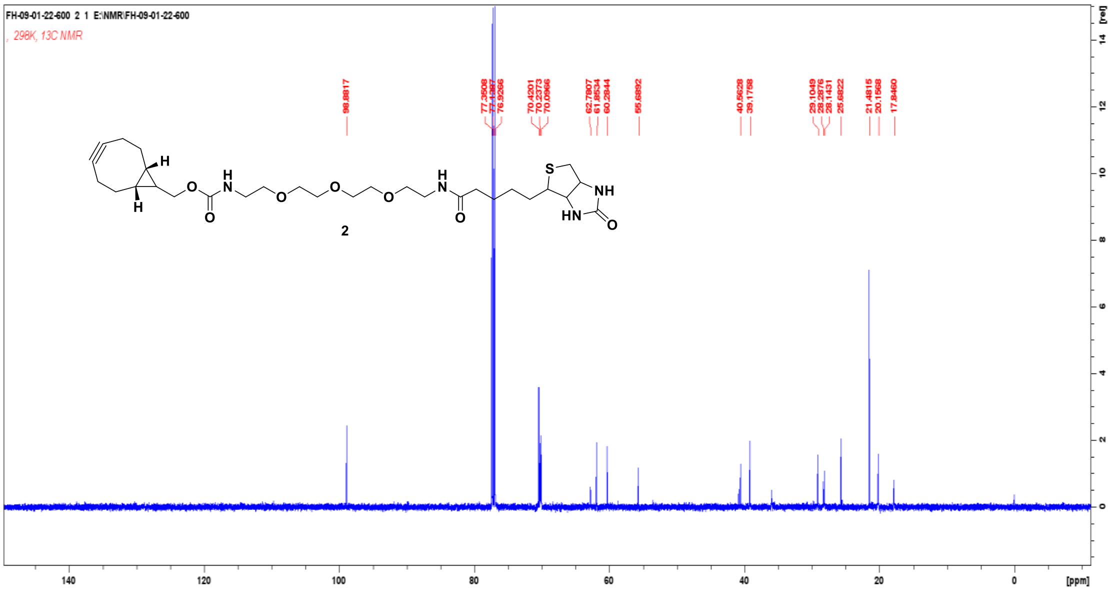

**Figure S4.** HRMS spectrum of compound **2**.

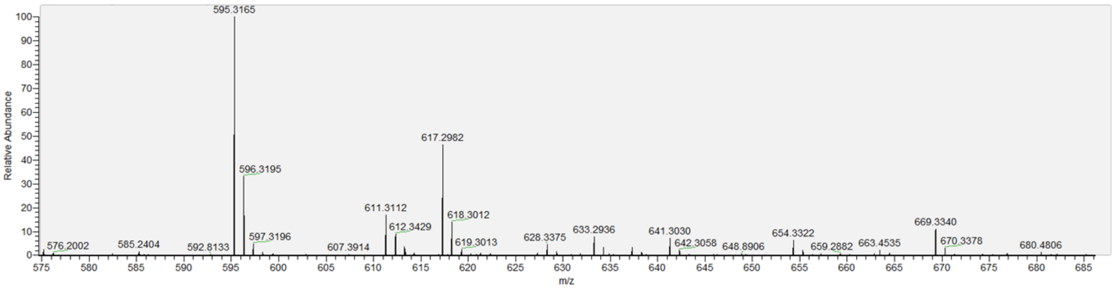

**Figure S5.** Structures and LC-MS analyses of the peptides used as substrates (A, B, C, D)and generated as products (E, F) modified by probes **1** and **2** in this study.

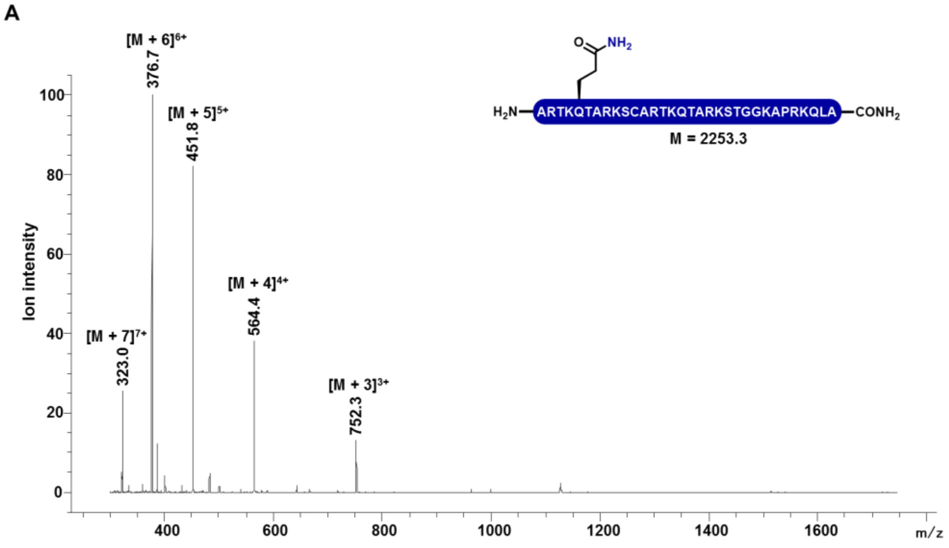


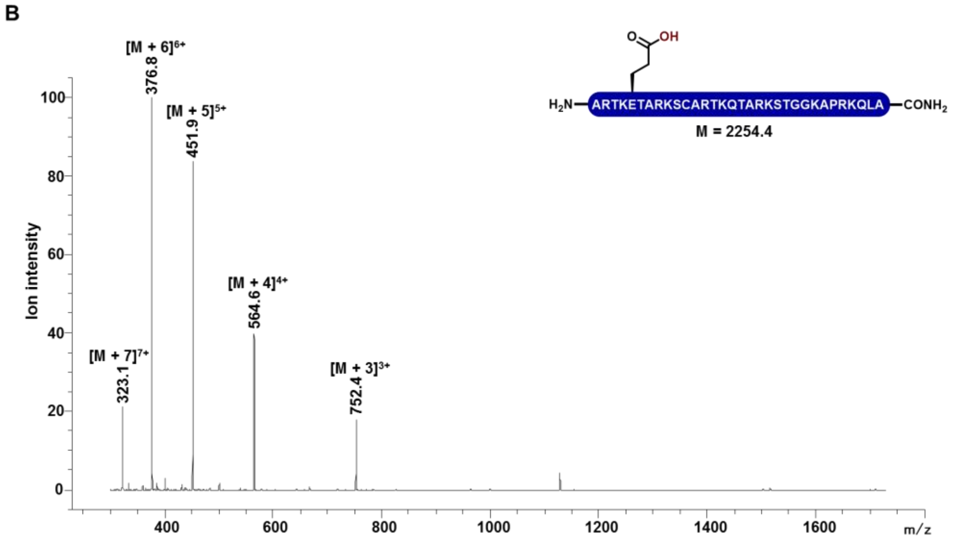


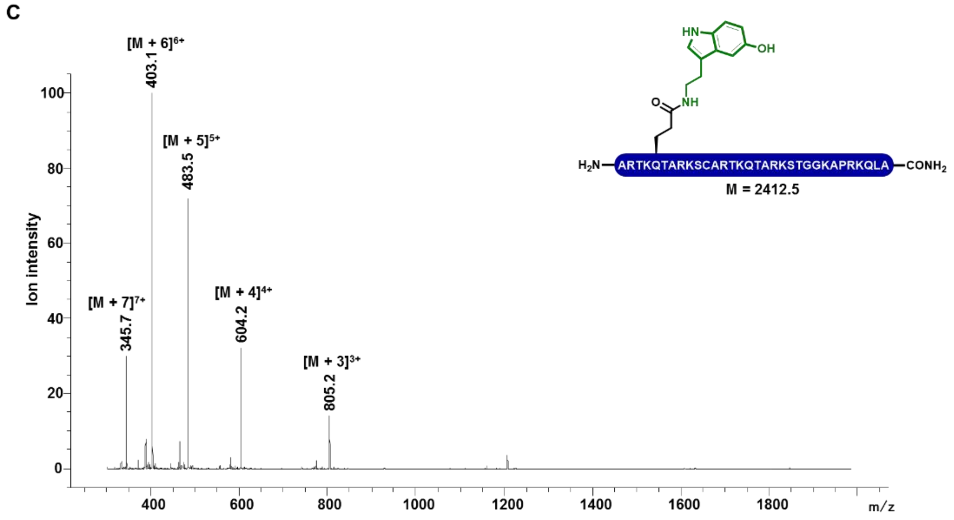

D

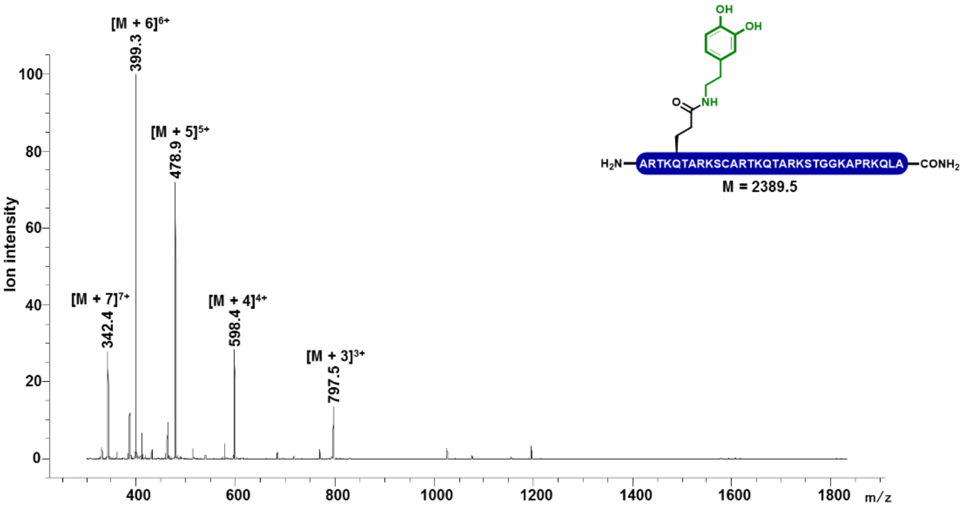
[M + 6]6+ 399.3
[M + 5]5+ 478.9
[M + 7]7+ 342.4
[M + 4]4+ 598.4
[M + 3]3+ 797.5
Ion intensity
m/z
H2N–ARTKQTARKSCARTKQTARKSTGGKAPRKQLA–CONH2
M = 2389.5

E

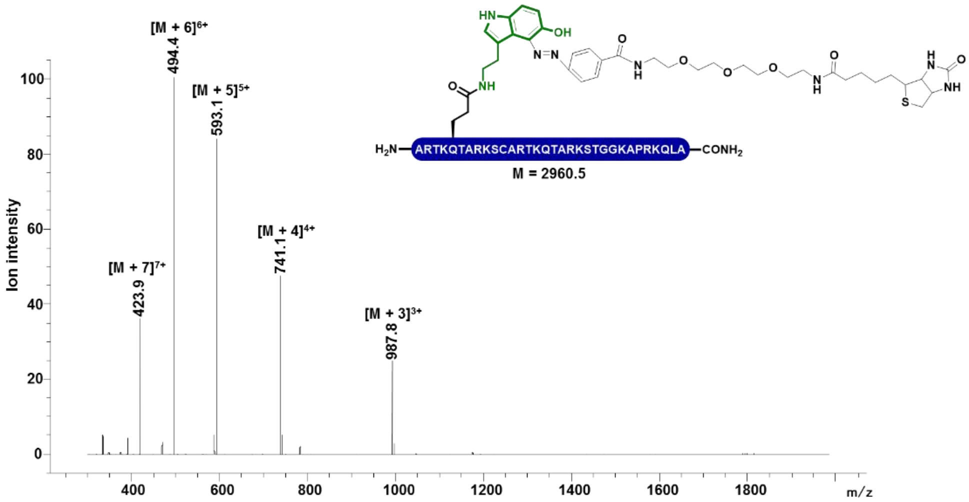
[M + 6]6+ 494.4
[M + 5]5+ 593.1
[M + 7]7+ 423.9
[M + 4]4+ 741.1
[M + 3]3+ 987.8
Ion intensity
m/z
H2N–ARTKQTARKSCARTKQTARKSTGGKAPRKQLA–CONH2
M = 2960.5

F

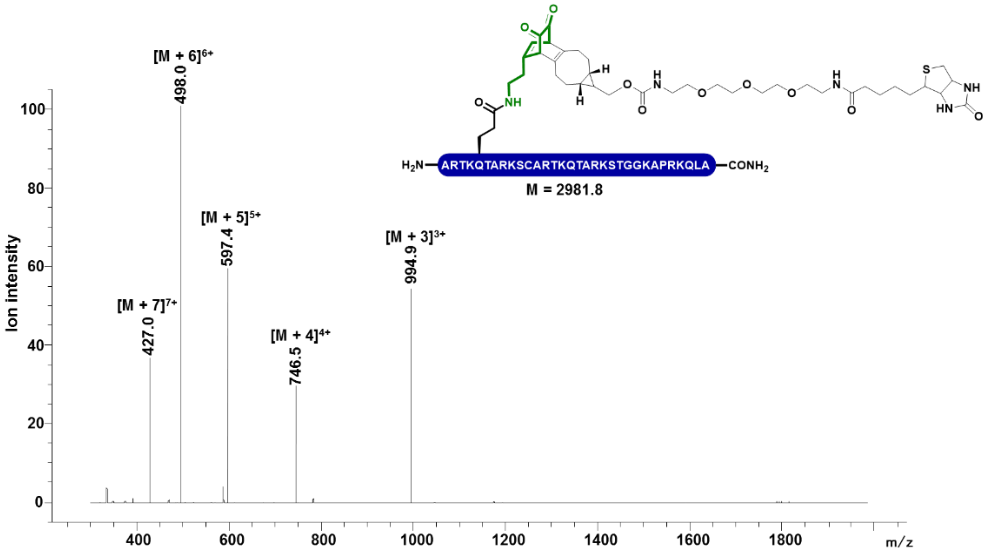
[M + 6]6+ 498.0
[M + 5]5+ 597.4
[M + 7]7+ 427.0
[M + 4]4+ 746.5
[M + 3]3+ 994.9
Ion intensity
m/z
H2N–ARTKQTARKSCARTKQTARKSTGGKAPRKQLA–CONH2
M = 2981.8

**Figure S6.** APAGE and in-gel imaging analysis of serotonin-modified nucleosomal arrays using probe **3** (A) and $Mg^{2+}$ precipitation analysis of the compaction states of modified arrays (B). Data are presented as mean values +/- SEM.

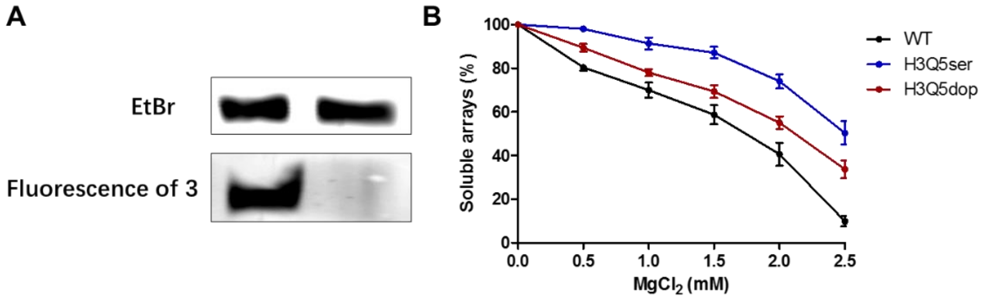
